# Enhanced phenylalanine biosynthesis amplifies light-stress-driven phenylpropanoid production in Arabidopsis

**DOI:** 10.64898/2026.02.16.706232

**Authors:** Rhowell Jr. N. Tiozon, Sara Christina Stolze, Anne Harzen, Hirofumi Nakagami, Hiroshi A. Maeda, Alisdair R. Fernie, Ryo Yokoyama

**Affiliations:** Max Planck Institute of Molecular Plant Physiology, Potsdam-Golm, Germany; Consumer-driven Grain Quality and Nutrition, Rice Breeding Innovation, International Rice Research Institute, Los Baños, Philippines; Max Planck Institute for Plant Breeding Research, Cologne, Germany; Department of Botany, University of Wisconsin-Madison, Madison, Wisconsin, USA; Division of Plant Science and Technology, University of Missouri, Columbia, Missouri, USA

## Abstract

Phenylpropanoids are L-phenylalanine (Phe)-derived specialized metabolites with crucial roles in plant stress adaptation. Among various abiotic stresses, high-light (HL) is a major threat that leads to oxidative damage and therefore upregulates the biosynthesis of antioxidant phenylpropanoids, including anthocyanins, in plants. Phenylpropanoid production is initiated by the conversion of Phe by phenylalanine ammonia-lyase (PAL) enzyme, which acts as the key gatekeeper controlling carbon influx from the Phe pool into phenylpropanoid biosynthesis. Despite the importance of Phe as a precursor for phenylpropanoid production, we have limited knowledge of how Phe precursor availability influences downstream pathways, particularly in the context of stress adaptation. To tackle this question, *Arabidopsis thaliana* wild-type and *production of anthocyanin pigment 1 dominant* (*pap1D*) mutant, a line genetically enhanced for anthocyanin production, were subjected to HL treatment, followed by proteome and metabolome analyses. Our multi-omics data indicated that Phe biosynthesis was insufficiently responsive to meet the increased precursor demand from the downstream phenylpropanoid pathway, likely making Phe availability a rate-limiting factor for HL-induced phenylpropanoid production. We then generated *pap1D* double mutants with Phe-overaccumulating mutants. Untargeted metabolomics revealed that enhanced Phe availability had little impact on phenylpropanoid accumulation under standard growth conditions but additively promoted the accumulation of specific phenylpropanoid intermediates and anthocyanin species under HL stress. This metabolic difference between light treatment and genotypes was correlated with the activation of PAL enzymatic activity. This study demonstrates that enhanced Phe biosynthesis amplifies HL-induced phenylpropanoid biosynthesis.

## Introduction

Phenylpropanoids constitute the largest and one of the most diverse classes of plant specialized metabolites, derived from the aromatic amino acid L-phenylalanine (Phe). This metabolic network gives rise to a diverse array of compounds, including flavonoids such as anthocyanins and flavonols, hydroxycinnamic acids, and lignin precursors, which collectively play central roles in plant growth, development, and environmental adaptation. Many flavonoids, particularly anthocyanins, function as photoprotective pigments, antioxidants, and signaling molecules, while hydroxycinnamate derivatives contribute to UV screening and redox homeostasis (Dixon and Paiva, 1995; Gould, 2004; Araguirang and Richter, 2022). In parallel, the phenylpropanoid pathway supplies monolignols for the biosynthesis of lignin, the primary cell wall component essential for vascular integrity and mechanical strength (Vogt, 2010; Vanholme *et al*., 2010; Dixon and Barros, 2019). Together, these metabolites enable plants to dynamically adjust both physiology and structure in response to fluctuating environmental conditions.

The biosynthesis of phenylpropanoids is initiated by the deamination of Phe through phenylalanine ammonia-lyase (PAL), linking primary amino acid metabolism to specialized metabolism (**Figure S1**) (Barros and Dixon, 2020). Following PAL-catalyzed deamination, cinnamic acid serves as a central branch point that is further modified by cinnamate 4-hydroxylase (C4H) and 4-coumarate:CoA ligase (4CL) to generate *p*-coumaroyl-CoA, a key intermediate at the interface of multiple phenylpropanoid branches toward the biosynthesis of hydroxycinnamate esters, monolignols for lignin formation. Entry into the anthocyanin branch is catalyzed by chalcone synthase (CHS), followed by chalcone isomerase (CHI), flavanone 3-hydroxylase (F3H), and downstream tailoring enzymes, including dihydroflavonol reductase (DFR) and anthocyanidin synthase (ANS), ultimately yielding diverse anthocyanin pigments through glycosylation and acylation reactions (**Figure S1**) (Vogt, 2010; Fraser and Chapple, 2011; Tohge *et al*., 2013). Over the past decades, extensive work has elucidated the transcriptional architecture governing phenylpropanoid biosynthesis, revealing a multilayered regulatory system involving MYB, bHLH, and WD40 transcription factors (Broun, 2005; Gonzalez *et al*., 2008; Hichri *et al*., 2011; Shin *et al*., 2015; Xie *et al*., 2016; Xing *et al*., 2024). A striking example is the isolation and characterization of the *production of anthocyanin pigment 1 dominant* (*pap1D*) mutant of *Arabidopsis thaliana*, in which activation tagging of the R2R3-MYB transcription factor PAP1 (MYB75) leads to massive and constitutive induction of phenylpropanoid biosynthetic genes and broad accumulation of flavonoids and lignin-related compounds across tissues and developmental stages (Borevitz *et al*., 2000; Tohge *et al*., 2005; Teng *et al*., 2005; Rowan *et al*., 2009; Shi and Xie, 2010). Under various environmental stresses, including high-light (HL), UV radiation, nutrient limitation, and pathogen challenge, this PAP1-mediated transcriptional network is particularly crucial for inducing expression of numerous biosynthetic genes, thereby enabling rapid production of stress-responsive phenylpropanoid compounds (Dixon and Paiva, 1995; Hichri *et al*., 2011; Dong and Lin, 2021; Li and Ahammed, 2023). Additionally, these findings establish transcriptional activation as a powerful approach for engineering phenylpropanoid accumulation, demonstrating that phenylpropanoid pathways can be genetically hyperactivated to the levels that are beyond their natural limits (Dixon *et al*., 1996; Broun, 2005; Grotewold, 2008; Zhu *et al*., 2020; Lee *et al*., 2024; Van Beirs *et al*., 2025).

In contrast to the well-characterized transcriptional control of phenylpropanoid biosynthesis, the upstream production of Phe is primarily regulated at the biochemical levels. Phe is synthesized via the shikimate pathway, which connects central carbon metabolism to the biosynthesis of three aromatic amino acids, Phe, L-tyrosine, and L-tryptophan (Tyr and Trp, respectively) (Tzin and Galili, 2010; Maeda and Dudareva, 2012; Lynch and Dudareva, 2020; El-Azaz and Maeda, 2025). The entry reaction of the shikimate pathway is catalyzed by 3-deoxy-D-*arabino*-heptulosonate-7-phosphate synthase (DAHP synthase or DHS), which condenses phospho*enol*pyruvate from glycolysis with D-erythrose-4-phosphate derived largely from the pentose phosphate pathway and the Calvin-Benson cycle in photosynthetic tissues (**Figure S1**) (Yokoyama *et al*., 2021; Yokoyama, Kleven, *et al*., 2022). Subsequent enzymatic steps in the shikimate pathway include 5-enolpyruvylshikimate-3-phosphate synthase (EPSPS), the target enzyme for glyphosate, and eventually generate chorismate, a common precursor for Phe/Tyr and Trp biosynthetic branches (Klee *et al*., 1987; Radwanski and Last, 1995; Tzin and Galili, 2010; Pollegioni *et al*., 2011; Maeda and Dudareva, 2012; El-Azaz and Maeda, 2025). In most vascular plants, chorismate mutase (CM) and prephenate aminotransferase dominantly produce arogenate, which is further converted into Phe and Tyr by arogenate dehydratase (ADT) and arogenate dehydrogenase (ADH or TyrA), respectively (**Figure S1**) (Jung *et al*., 1986; Romero *et al*., 1995; Cho *et al*., 2007; Tzin and Galili, 2010; Maeda *et al*., 2010; Maeda and Dudareva, 2012; de Oliveira *et al*., 2019). Phe is utilized not only for protein synthesis but also serves as an indispensable precursor for a wide range of specialized metabolites, most notably phenylpropanoids, as well as other Phe-derived compounds such as phenylpyruvate, which is produced directly from Phe by aromatic aminotransferases independently of the phenylpropanoid pathway (**Figure S1**) (Koper *et al*., 2022; Koper *et al*., 2025). Because aromatic amino acid homeostasis is critical for both plant growth and natural product biosynthesis, the aromatic amino acid biosynthetic pathway is tightly regulated. Key enzymatic steps, including DHS, CM, ADT, and TyrA, are controlled by negative feedback inhibition from downstream aromatic amino acids and related intermediates (Maeda and Dudareva, 2012; Yokoyama, 2024; El-Azaz and Maeda, 2025). Introducing genetic mutations that deregulate these negative feedback inhibitions leads to an increase in aromatic amino acid accumulation, including *suppressor of tyra* (*sota*) mutations that deregulate DHS feedback inhibition and boost aromatic amino acid production (Yokoyama, de Oliveira, *et al*., 2022). Recently, enzyme isoforms with relaxed feedback inhibition have been identified in specific plant lineages that support high production of aromatic amino acid-derived metabolites, highlighting the key role of amino acid precursor supply in expanding chemical diversity during plant evolution (Maeda, 2019a; Yokoyama, 2024). Compared with such long-term evolutionary adjustment, whether plants have a mechanism to adjust precursor supply in response to shorter-term changes, such as abiotic stresses in which plants induce the production of Phe-derived anti-stress metabolites, remains poorly understood. The lack of mechanistic understanding of the coordination between Phe precursor supply and downstream phenylpropanoid pathway has also been a major bottleneck in improving metabolic engineering of Phe-derived specialized metabolites.

Here, we address this unresolved fundamental knowledge gap by testing the hypothesis that Phe availability constitutes a metabolic bottleneck for HL-induced phenylpropanoid biosynthesis. Specifically, we ask whether enhancing Phe biosynthetic capacity is sufficient to amplify phenylpropanoid accumulation under HL conditions. By disentangling precursor supply from pathway induction, this work provides new insight into how primary and specialized metabolism are coordinately regulated during environmental stress and reveals Phe precursor availability as a critical determinant of phenylpropanoid production under abiotic stress conditions.

## Results

### Anthocyanin overproduction led to a Phe precursor shortage

To examine how Phe content alters upon activation of Phe-derived specialized metabolism genetically and environmentally, Col-0 and *pap1D* plants were grown under standard short-day light condition (8-h light, 100 µmol m^-2^ s^-1^) and exposed to continuous HL condition at 600 µmol m^-2^ s^-1^ for 2 days (**Figure 1A**). Before HL treatment, *pap1D* accumulated 2.7 times more anthocyanins than Col-0 (**Figure 1B**), consistent with prior studies (Borevitz *et al*., 2000). Two-day HL exposure led to 7.8- and 7.0-fold increases in anthocyanin content in Col-0 and *pap1D*, respectively, resulting in 18.7-fold higher anthocyanin accumulation in HL-treated *pap1D* plants compared with untreated Col-0 plants (**Figure 1B**). To better resolve individual phenylpropanoid compounds, targeted liquid chromatography-mass spectrometry (LC-MS) metabolomic analysis was performed using the same plant samples. In Arabidopsis, eleven types of structurally distinct cyanidin-derived anthocyanin species have been reported previously (Tohge *et al*., 2005; Yonekura-Sakakibara *et al*., 2012). Among the nine anthocyanin species detected in this analysis, most showed higher levels in *pap1D* than in Col-0 both before and after HL treatment, and their contents increased in response to HL (**Figure S2**). As an exception, anthocyanin A6 (Cyanidin 3-*O*-[2′′-*O*-(xylosyl)-6′′-*O*-(*p*-*O*-(glucosyl)-*p*-coumaroyl) glucoside] 5-*O*-glucoside) accumulated to a lower level in *pap1D* than in Col-0 after HL exposure (**Figure S2**). HL treatment also elevated the levels of several other phenylpropanoids, including flavonols, in both genotypes (**Figure S3**). Like anthocyanin A6, kaempferol- or apigenin-conjugated flavonols, but not quercetin-conjugated ones, were accumulated to lower levels in *pap1D* than in Col-0 after HL treatment (**Figure S3**). Overall, while both HL exposure and the *pap1D* mutation individually stimulated phenylpropanoid biosynthesis, their combination further enhanced the accumulation of anthocyanins and other phenylpropanoids.

**Figure 1.**
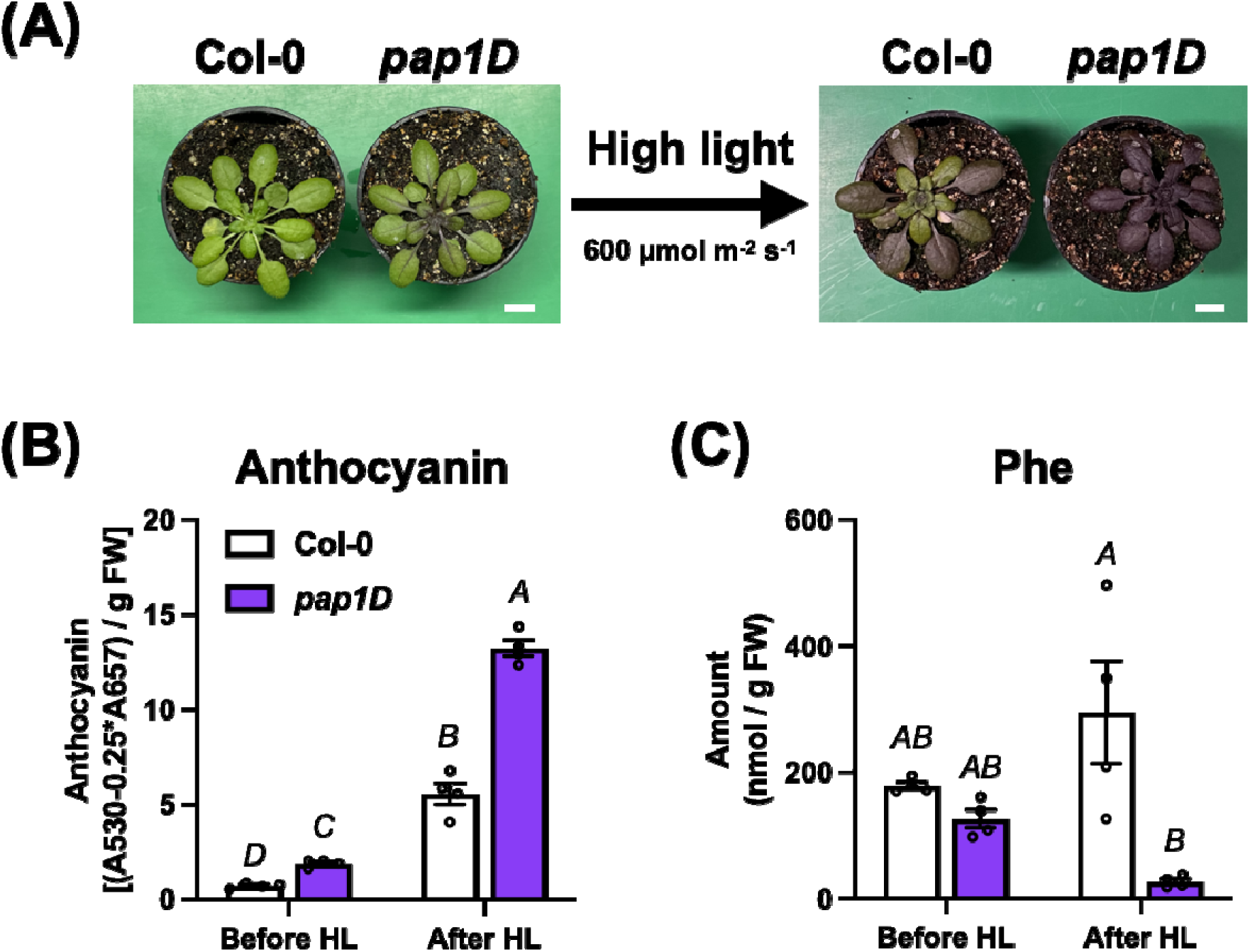
Anthocyanin overproduction resulted in Phe depletion. **(A)** Plant pictures of Col-0 and *pap1D* before and after HL exposure (600 µmol m^-2^ s^-1^ for 2 days). Bar = 1 cm. **(B and C)** Metabolite levels of anthocyanin (B) and Phe (C) in Col-0 and *pap1D* before and after HL. Different letters indicate statistically significant differences (two-way ANOVA with Tukey-Kramer test, *P* < 0.05). Data are means ± SEM (*n* = 4).

Phe content was also quantified in the same samples to examine how Phe availability was changed in response to upregulation of phenylpropanoid production during HL stress. Before HL treatment, the Phe levels were comparable between Col-0 and *pap1D*. After HL, Phe content remained statistically unchanged in Col-0 but was significantly lower in *pap1D* than in Col-0 (**Figure 1C**). This result indicated that overproduction of anthocyanins and other Phe-derived phenylpropanoids led to Phe depletion, likely due to increased metabolic demand of Phe as a precursor for phenylpropanoid synthesis under HL condition. Because the *pap1D* mutation alone did not affect Phe levels before HL exposure, HL-induced activation of phenylpropanoid metabolism appears to have a more pronounced impact on Phe precursor availability than the *pap1D* mutation itself.

### Phe biosynthesis was less responsive to HL at the proteome level than its downstream pathways

Abiotic stresses, including HL, often induce dynamic remodeling of the proteome, leading to the acclimation under adverse environmental conditions (Giacomelli *et al*., 2006; Miller *et al*., 2017; Watson *et al*., 2018; Kosová *et al*., 2018). While responses of specialized metabolism to HL have been extensively characterized, much less is known about how upstream primary metabolism, including Phe metabolism, responds to HL at the proteomic level. To address this gap, a comparative proteome analysis was performed using Col-0 and *pap1D* before and after HL treatment. In Col-0 and *pap1D*, HL exposure upregulated 211 and 530 proteins and downregulated 229 and 577 proteins, respectively, suggesting broader impacts on the proteome in *pap1D* than in Col-0 (**Figure 2A and 2B**). Well-established HL marker proteins, including Early Light-Induced Protein 1 (ELIP1, AT3G22840) (Hutin *et al*., 2003; Giacomelli *et al*., 2006), were significantly upregulated by HL (**Figure 2C and 2D**), confirming that our HL treatment was effective. The gene set enrichment analysis (GSEA) revealed that secondary (specialized) metabolic pathways, including those related to flavonoid biosynthesis, were significantly enriched among proteins whose abundance increased under HL stress (**Figure 2E**). Consistent with this enrichment, several core anthocyanin biosynthetic enzymes, including CHS, DFR, and UDP-flavonoid 3-*O*-glucosyltransferase (3GT), were upregulated following HL treatment (**Figures 2C and 2E**). This finding indicates that proteome-level activation of photoprotective phenylpropanoid metabolism constitutes a major component of the Arabidopsis acclimation response to HL at the proteome level. In contrast, our GSEA failed to identify any GO terms associated with shikimate or Phe biosynthesis among HL-responsive proteins (**Figure 2E**). This absence of enrichment indicates that, unlike downstream phenylpropanoid metabolism, Phe biosynthesis is not coordinately upregulated at the proteome level in response to HL stress.

**Figure 2.**
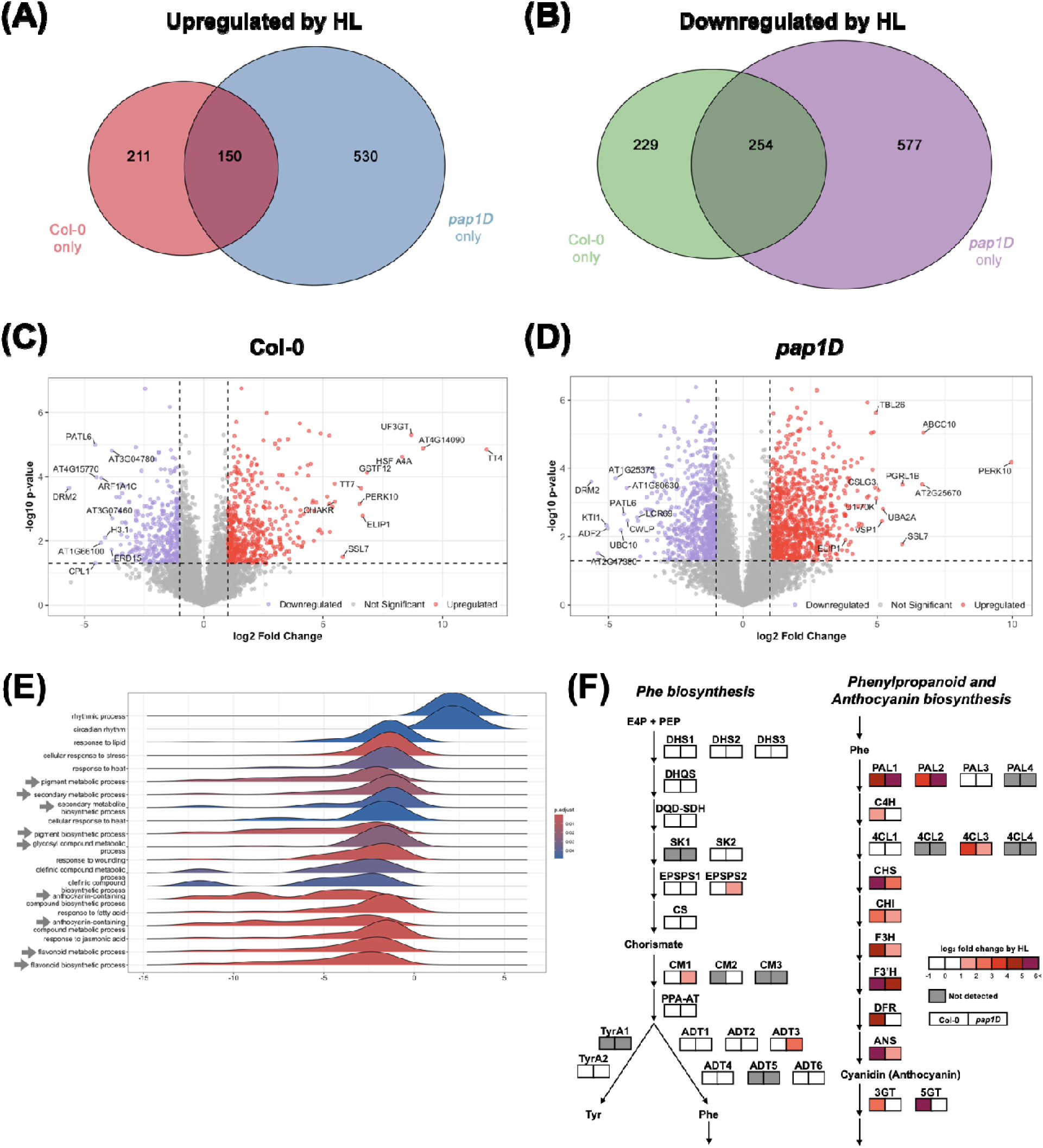
Phe biosynthesis was less responsive to HL stress than the phenylpropanoid pathway. **(A, B)** Venn diagrams of the numbers of proteins upregulated (A) and downregulated (B) by HL in Col-0 and *pap1D*. **(C, D)** Volcano plots of proteins upregulated (red) and downregulated (blue) by HL in Col-0 (C) and *pap1D* (D). **(E)** Ridge plots depict significantly enriched GO biological process terms based on changes in protein abundance that are upregulated in Col-0 and *pap1D* under HL. Color scale denotes adjusted *p*-values. GO terms related to phenylpropanoid metabolism were pointed by gray arrows **(F)** Fold changes in the levels of proteins participating in Phe biosynthesis (left) and phenylpropanoid and anthocyanin biosynthesis (right) after HL exposure. The left and right boxes of each enzymatic step represent fold changes in Col-0 and *pap1D*, respectively.

To further visualize differences in proteome responses between Phe biosynthesis and the downstream pathway, we mapped the HL-induced fold changes in protein abundance onto the Phe and anthocyanin biosynthetic pathways. Consistent with the GSEA results, most enzymes involved in the early phenylpropanoid pathway and anthocyanin biosynthesis, including inducible PAL1 and PAL2 isoforms (Raes *et al*., 2003; Rohde *et al*., 2004; Barros and Dixon, 2020), were upregulated in both Col-0 and *pap1D* by HL (**Figure 2F**). In contrast, the protein levels of enzymes in the upstream shikimate and Phe biosynthetic pathways were largely unchanged, except for a slight increase in EPSPS2, CM1, and ADT3 in *pap1D* (**Figure 2F**). Collectively, these results demonstrate that Phe biosynthesis is considerably less responsive to HL exposure than its downstream phenylpropanoid and anthocyanin biosynthesis. This imbalance of HL responsiveness between Phe biosynthesis and the downstream pathway may contribute to a shortage of Phe precursors under HL-induced activation of phenylpropanoid metabolism (**Figures 1C and 1D**).

### HL had a broader impact on the metabolome than on the *pap1D* mutation

These multi-omics analyses of *pap1D* suggested that Phe availability was a bottleneck to produce anthocyanins and other phenylpropanoids under HL conditions. To test this hypothesis, we artificially enhanced Phe availability in *pap1D* and analyzed its impacts on downstream phenylpropanoid biosynthesis. The *pap1D* mutant was crossed with two independent allelic *sota* mutants, *sotaB4* and *sotaA4*, which hyperaccumulate Phe due to the deregulation of DHS negative feedback inhibition by point mutations on the *DHS1* and *DHS2* genes, respectively (Yokoyama *et al*., 2021; Yokoyama, de Oliveira, *et al*., 2022). The resulting double mutants, *pap1D sotaB4* and *pap1D sotaA4*, along with their corresponding single mutants were then subjected to the same 2-day HL treatment (**Figure 3A**). Consistent with the prior publication (Yokoyama, de Oliveira, *et al*., 2022), neither *sotaB4* nor *sotaA4* mutant showed elevated anthocyanin levels before HL exposure (**Figure 3B**). In the *pap1D sotaB4* and *pap1D sotaA4* double mutants, the amounts of anthocyanins were higher than the respective *sota* single mutants, but at the same levels as the *pap1D* single mutant (**Figure 3B**), indicating that enhanced Phe production does not promote anthocyanin content even in the *pap1D* mutant background under the standard growth condition. To test whether elevated Phe availability altered HL-driven phenylpropanoid production, we subjected the same Arabidopsis plants to 2-day continuous HL treatment. While the anthocyanin amount was elevated by HL stress in all the genotypes, *sotaB4* and *pap1D sotaA4* showed a slightly but statistically higher levels of anthocyanin than Col-0 and *pap1D*, respectively (**Figure 3B**). This result suggests that elevated Phe production by the *sota* mutations might have some impact on anthocyanin production under HL condition. However, absorbance-based anthocyanin quantification may be affected by interference from co-extracted non-anthocyanin chromophores that contribute to the absorbance signal and mask actual variation in anthocyanin and other phenylpropanoid accumulation (Wrolstad *et al*., 2005; Welch *et al*., 2008).

**Figure 3.**
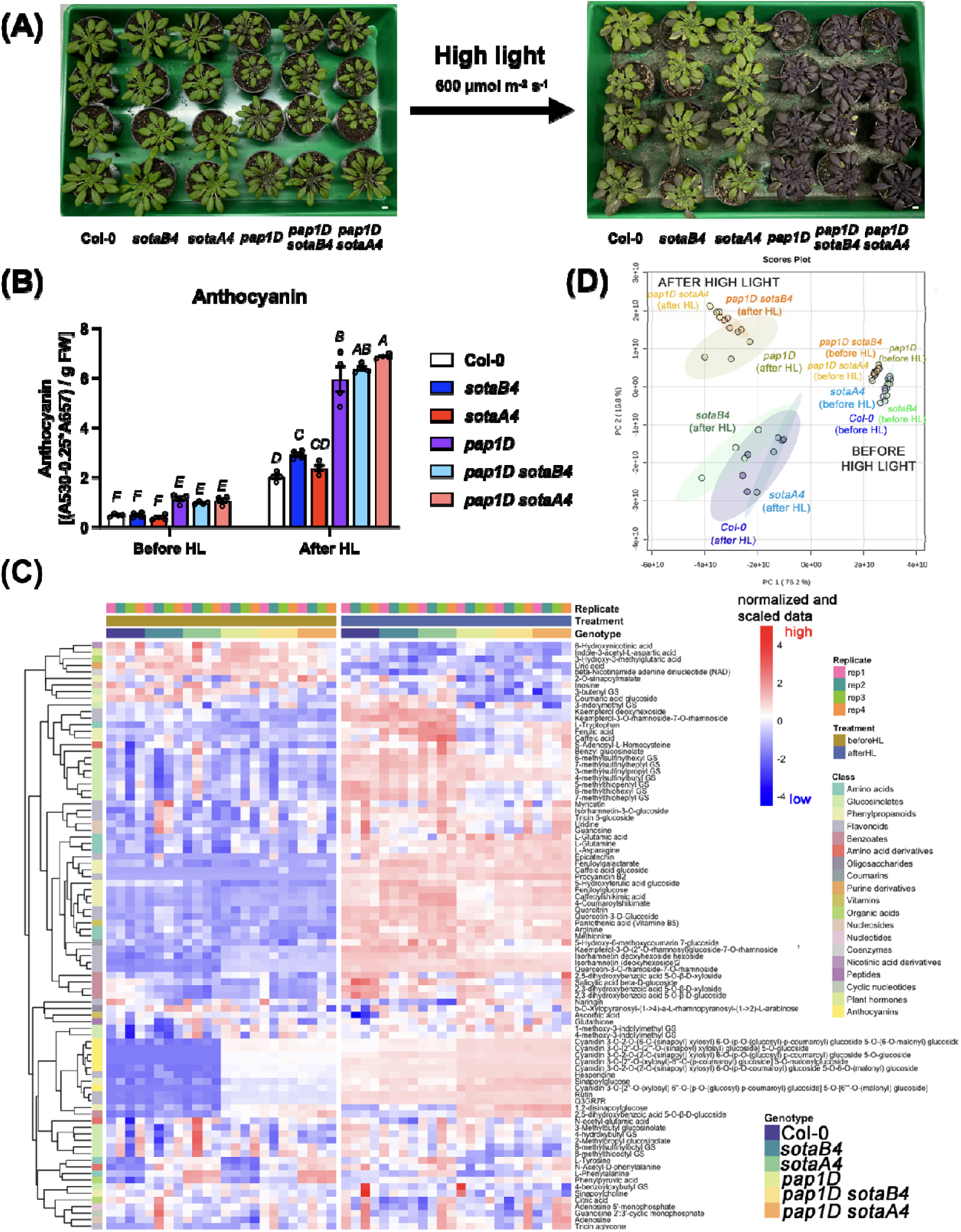
Untargeted metabolomics revealed the broader impact of HL exposure on the metabolome. **(A)** Plant pictures of Col-0, *sotaB4*, *sotaA4*, *pap1D*, *pap1D sotaB4*, and *pap1D sotaA4* before and after HL exposure (600 µmol m^-2^ s^-1^ for 2 days). Bar = 1 cm. **(B)** Anthocyanin contents of Col-0, *sotaB4*, *sotaA4*, *pap1D*, *pap1D sotaB4*, and *pap1D sotaA4* before and after HL exposure. Different letters indicate statistically significant differences (two-way ANOVA with Tukey-Kramer test, *P* < 0.05). Data are means ± SEM (*n* = 4). **(C)** Hierarchical clustering heatmap of normalized and scaled metabolite abundances for 89 annotated metabolites. Columns represent biological replicates grouped by genotype and treatment, and rows represent individual metabolites clustered based on abundance patterns. Metabolite classes are indicated by color bars. **(D)** Principal component analysis (PCA) of untargeted metabolomic profiles derived from Arabidopsis Col-0, *sotaB4*, *sotaA4*, *pap1D*, *pap1D sotaB4*, and *pap1D sotaA4* plants before and after HL treatment. Ellipses represent 95% confidence intervals. PERMANOVA analysis confirms a significant treatment-driven separation of metabolite profiles (F = 127.13, R² = 0.9749, *p* = 0.001).

To examine the impacts of the *sota* mutations on the metabolome more comprehensively and quantitatively, the same plant samples were subjected to untargeted metabolomics using LC-MS/MS in negative ion mode. In this study, we annotated 89 metabolites based on standards, and/or MS/MS data from previous studies (**Supplemental Data Set 8**), and this number was close to that previously reported in Arabidopsis metabolomics studies (Tohge *et al*., 2016; Wu *et al*., 2018; Naake *et al*., 2024), suggesting that our annotation approach was robust. To visualize global changes in metabolite abundance across genotypes and treatments, the normalized and scaled metabolite profiles were subjected to hierarchical clustering and displayed as a heatmap (**Figure 3C**). The heatmap revealed a pronounced and coordinated shift in the metabolome upon HL exposure across all genotypes, with samples collected after HL forming a distinct cluster separate from those harvested before HL. This HL-driven reprogramming was characterized by increased accumulation of phenylpropanoids, including flavonols, anthocyanins, coumarins, and hydroxycinnamate derivatives, alongside changes in amino acids and related specialized metabolites, whereas other classes of specialized metabolites, such as glucosinolates, were downregulated by HL exposure (**Figures 3C and S4**). In contrast, genotype-dependent differences were comparatively subtle under control conditions, while modest separation was still observed between lines with and without the *pap1D* mutation prior to HL treatment. Notably, after HL exposure, *pap1D* and *pap1D sota* lines exhibited broadly similar metabolite accumulation patterns, indicating that HL-induced metabolic remodeling largely overrides genotype-specific baseline differences arising from the *pap1D* mutation or elevated Phe availability. Concurrently, a principal component analysis (PCA) based on specialized metabolite profiles revealed a clear separation between samples collected before and after HL treatment along the first principal component (PC1), which explained 76.2% of the total variance, indicating that HL exposure was the dominant factor shaping the metabolome. In contrast, genotype-dependent differences were captured mainly along PC2 (16.8% of the variance), with the *pap1D* and *pap1D sota* lines clustering more closely after HL treatment (**Figure 3D**). Together, these results demonstrate that HL treatment exerts a stronger effect on global specialized metabolite composition than the genetic backgrounds.

### Enhancing Phe biosynthesis additively promoted HL-induced phenylpropanoid production

To more closely investigate how elevated Phe precursor availability influences individual downstream pathways, we examined the levels of Phe and specific Phe-derived specialized metabolites across genotypes and under HL stress treatment. As reported previously (Yokoyama, de Oliveira, *et al*., 2022), the *sotaA4* allele exhibited the highest steady-state levels of all three aromatic amino acids, Phe, Tyr, and Trp, followed by *sotaB4*, which also showed significantly elevated aromatic amino acid levels (**Figures 4 and S5**). In the *pap1D sotaB4* and *pap1D sotaA4* double mutants, the amounts of Phe and the other aromatic amino acids were elevated at the same levels as the *sotaB4* and *sotaA4* single mutants, respectively (**Figures 4 and S5**). Consistent with the previous report (Yokoyama, de Oliveira, *et al*., 2022), the *sota* mutants exhibited the elevated levels of phenylpyruvate (**Figure 4**), which is directly produced from Phe by aromatic aminotransferases independently of the phenylpropanoid pathway (**Figure S1**). The *pap1D sota* double mutants also show the similar levels of phenylpyruvate to the *sota* single mutants. In contrast, the impact of enhanced Phe biosynthesis on the phenylpropanoid and anthocyanin pathways was limited (**Figure 4**). General phenylpropanoid intermediates, including 4-coumaroylshikimate, caffeoylshikimate, ferulic acid, and caffeic acid, and major flavonols, such as quercetin-3-*O*-(2"-*O*-rhamnosyl) glucoside-7-*O*-rhamnoside and kaempferol-3-*O*-(2"-*O*-rhamnosyl) glucoside-7-*O*-rhamnoside (Q3GR7R and K3GR7R, respectively), were accumulated at similar levels among all genotypes before HL treatment (**Figure 4**). While mutants harboring the *pap1D* background accumulated elevated levels of anthocyanin (e.g., anthocyanin A9, A10, and A11, among the most abundant anthocyanin species) and procyanidin B2, the *sota* single and double mutants had the same levels of anthocyanins as Col-0 and *pap1D* (**Figure 4**). Collectively, these results suggest that enhanced Phe availability does not promote phenylpropanoid or anthocyanin production in either the Col-0 or *pap1D* background before HL exposure.

**Figure 4.**
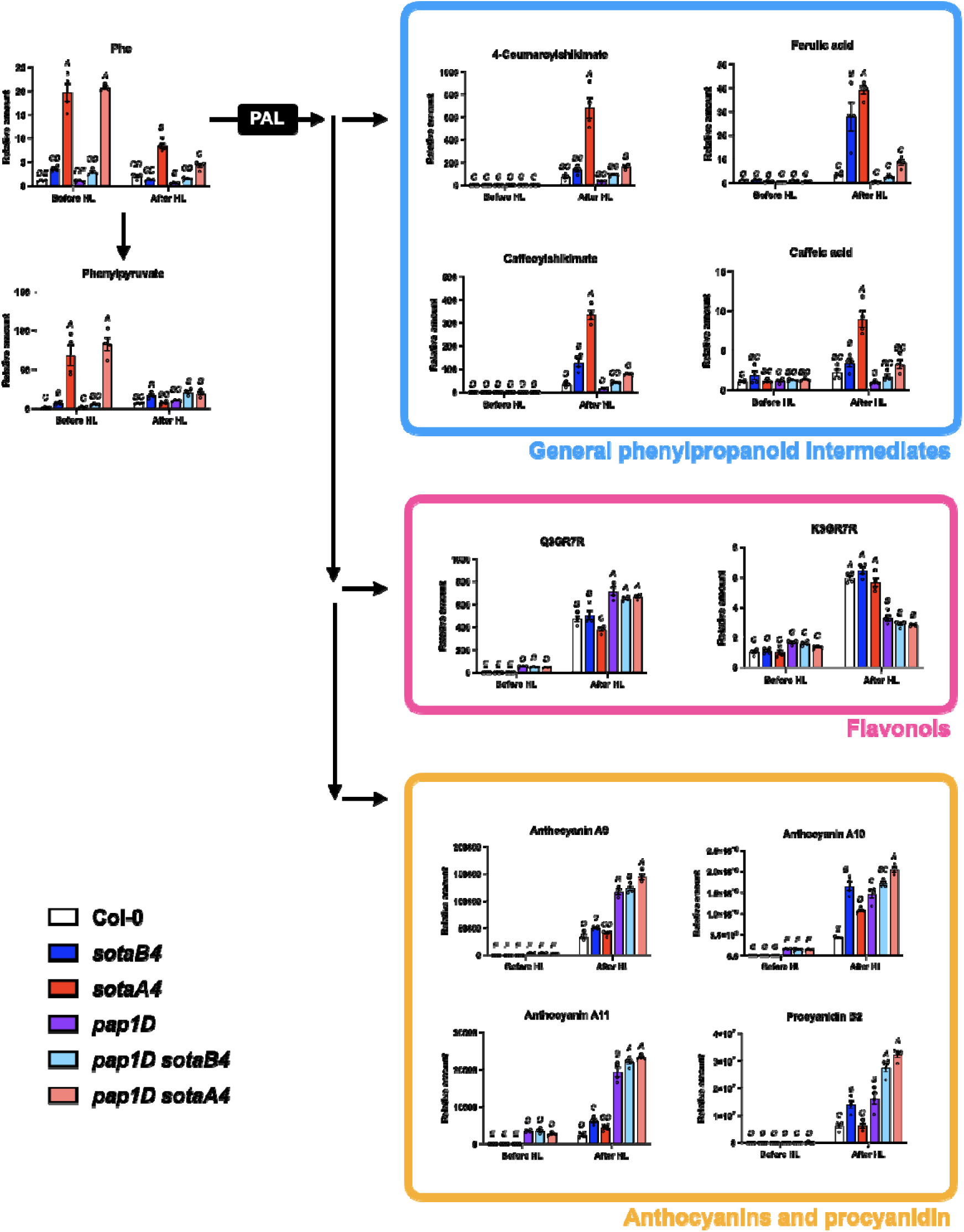
Increased Phe precursor availability additively promoted the accumulation of some phenylpropanoid compounds under HL stress. The levels of selected metabolites of Col-0, *sotaB4*, *sotaA4*, *pap1D*, *pap1D sotaB4*, and *pap1D sotaA4* before and after HL exposure. Different letters indicate statistically significant differences (two-way ANOVA with Tukey-Kramer test, *P* < 0.05). Data are means ± SEM (*n* = 4).

Under HL stress, the effect of the *sota* mutations on phenylpropanoid production became apparent. After HL exposure, Phe levels were markedly reduced in the *sotaA4* and *pap1D sotaA4* mutants, accompanied by a decrease in phenylpyruvate levels (**Figure 4**). In contrast, the levels of Tyr or Trp did not change as drastically as Phe after HL stress (**Figure S5**). Strikingly, the *sotaA4* mutant exhibited the highest accumulation of the general phenylpropanoid intermediates: ferulic acid, caffeic acid, caffeoylshikimate, and 4-coumaroylshikimate. The *pap1D sota* double mutants also displayed a similar pattern, accumulating higher levels of early phenylpropanoid intermediates than the *pap1D* single mutant (**Figure 4**). However, in plants carrying the *pap1D* mutant background, these intermediates were less abundant than in their respective Col-0 background lines (**Figure 4**), probably due to more dominant carbon utilization for their downstream anthocyanin pathway. The levels of some anthocyanin species and procyanidin B2 after HL exposure were higher in the *sota* mutants than in Col-0, and they were further increased in the *pap1D sota* double mutants relative to the *pap1D* single mutant (**Figure 4**). These results suggest that increased Phe levels contribute additively to the accumulation of certain anthocyanin species under HL conditions. The levels of kaempferol- or quercetin-conjugated flavonols were not enhanced under HL stress in mutants harboring the *sota* mutant background (**Figure 4**). In contrast to the HL-induced upregulation of phenylpropanoid and anthocyanin biosynthesis, the level of phenylpyruvate decreased under HL, particularly in *sotaA4* and *pap1D sotaA4*, both of which exhibited the highest Phe accumulation before HL exposure (**Figure 4**). This relationship between the phenylpropanoid and phenylpyruvate pathways implies that HL stress may channel carbon from the Phe pool preferentially toward the phenylpropanoid branch rather than the phenylpyruvate pathway.

### Impact of enhanced Phe supply on phenylpropanoid production positively correlated with PAL activity

The metabolomics data before and after HL exposure suggested that the HL treatment upregulated the activity of PAL enzymes, which catalyze the committed step of the phenylpropanoid pathway. To test this hypothesis, we measured PAL enzymatic activity using the same samples used for metabolomics analysis above. Before HL, PAL activity was comparable between Col-0 and the *sota* mutants. The presence of the *pap1D* activation tag, i.e., *pap1D* and *pap1D sota* mutants, showed elevated PAL activity, in agreement with the previous publication reporting that *pap1D* displayed >2-fold higher PAL activity than Col-0 under standard growth condition (**Figure 5**) (Borevitz *et al*., 2000). Notably, HL treatment dramatically enhanced PAL activity in all genotypes, with the *pap1D* and *pap1D sota* mutants showing the highest PAL activity (**Figure 5**). These results showed that the HL treatment can increase the PAL activity at the committed step of the phenylpropanoid biosynthesis. Given that the impact of the *sota* mutations on phenylpropanoid production became apparent only after HL exposure (**Figures 3B and 4**), this HL-induced PAL activity may be able to release the bottleneck and direct the accumulated Phe, caused by the *sota* mutations, towards the downstream phenylpropanoid production.

**Figure 5.**
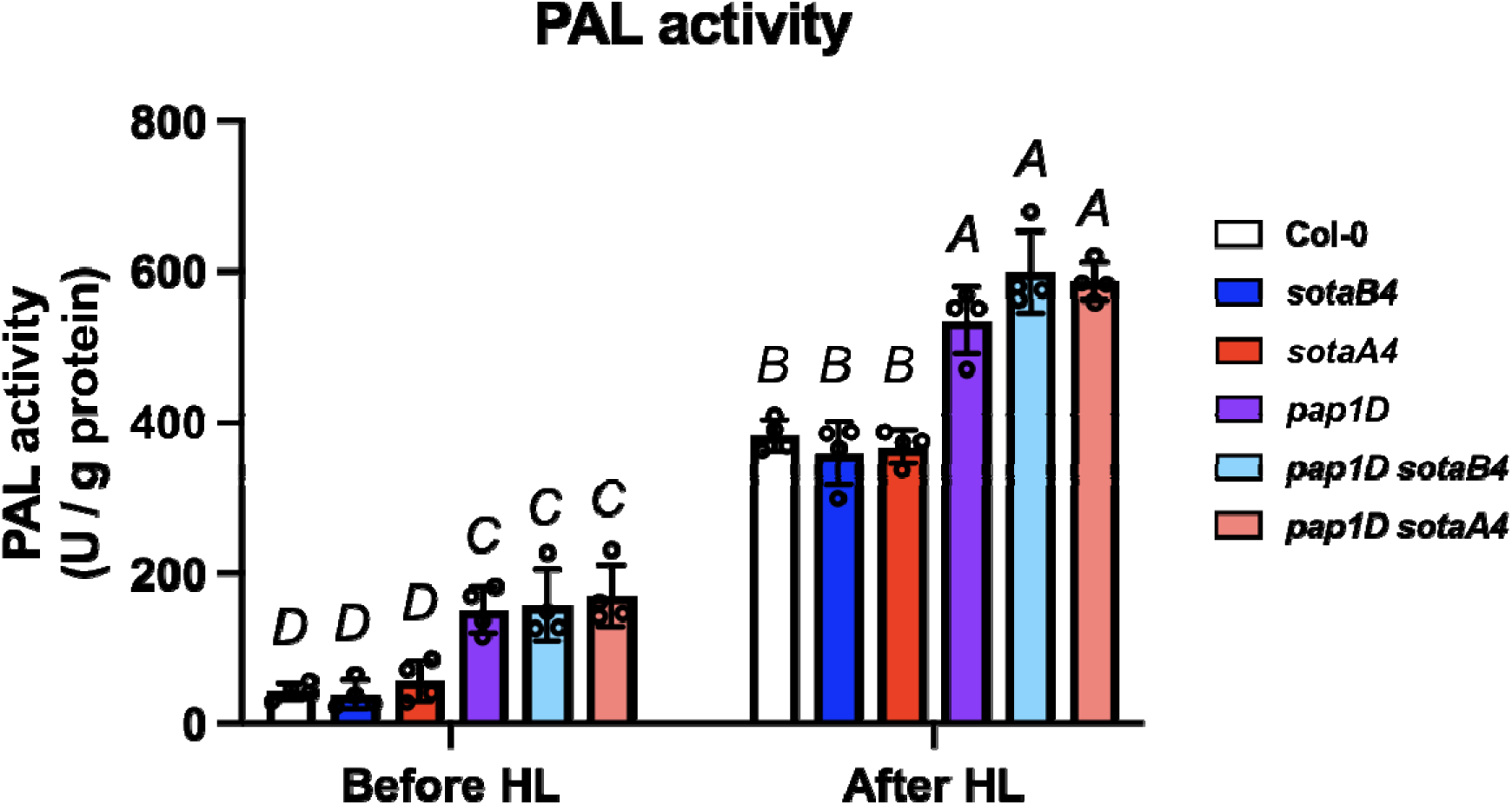
PAL enzymatic activity correlates with the efficiency of Phe utilization toward phenylpropanoid biosynthesis under HL stress. The PAL enzyme activity isolated from Col-0, *sotaB4*, *sotaA4*, *pap1D*, *pap1D sotaB4*, and *pap1D sotaA4* before and after HL exposure. Different letters indicate statistically significant differences (two-way ANOVA with Tukey-Kramer test, *P* < 0.05). Data are means ± SEM (*n* = 4).

## Discussion

### Phe precursor availability as a conditional bottleneck in high-stress-driven phenylpropanoid production

Focusing on anthocyanins, this study examines how the availability of Phe precursors influences downstream Phe-derived specialized metabolic pathways in Arabidopsis. We demonstrate that increased Phe availability has a limited impact on phenylpropanoid production under standard growth conditions but markedly boosts phenylpropanoid biosynthesis under HL stress (**Figures 3 and 4**). These findings suggest that accumulating excess Phe serves as an effective strategy to be prepared for future fluctuations in light intensity, enabling rapid induction of phenylpropanoid pathways when physiological demand increases.

The relationship between precursor supply and product accumulation appears complex, as evidenced by previous metabolic engineering studies in Arabidopsis. Heterologous expression of bacterial deregulated AroG (DHS) enzyme and bi-functional CM/prephenate dehydratase enzyme resulted in approximately 2.5-7.5-fold and 4-10-fold increase in free Phe content in 10-day-old Arabidopsis seedlings, respectively, accompanied by upregulated production of some phenylpropanoids, such as sinapyl alcohol, coniferin, and kaempferol deoxyhexose (Tzin *et al*., 2009; Tzin *et al*., 2012). Similarly, a deregulated ADT2 point mutation increased Phe production by approximately 70-fold, leading to a 2-fold elevation in kaempferol- and quercetin-conjugated flavonoids (Huang *et al*., 2010). In contrast, more than 20-fold upregulation of Phe production in the *sota* mutants does not enhance phenylpropanoid production under standard growth conditions (Yokoyama, de Oliveira, *et al*., 2022). Together, these studies demonstrate that increasing Phe abundance does not necessarily lead to proportional enhancement of phenylpropanoid production, implying that additional regulatory factors are required to effectively transmit changes in Phe availability to downstream phenylpropanoid biosynthesis. Although more direct genetic or pharmacological manipulation of PAL activity, such as *PAL* overexpression in the *sota* mutant background, will be required to establish the causality, our study indicates that PAL activation may be one of the critical factors determining whether enhanced Phe availability leads to increased phenylpropanoid production (**Figure 5**). We observed a positive correlation between HL-induced PAL upregulation and the additive effect of Phe overaccumulation on downstream metabolites. In other words, under standard growth conditions, basal PAL activity may impose a strict "gatekeeper" limitation on metabolic flux. HL exposure likely relieves this bottleneck by elevating PAL activity, thereby unlocking the latent capacity of the expanded Phe pool to drive phenylpropanoid biosynthesis. Phe precursor availability therefore becomes a conditional bottleneck under HL stress. The observation that some earlier studies reported increased phenylpropanoid production alongside elevated Phe levels may be explained by higher basal PAL activity associated with the specific environmental contexts or developmental stages used in those experiments.

Meanwhile, we cannot eliminate the possibility that other factors influence the conversion efficiency of excess Phe molecules into phenylpropanoids during HL stress. For example, it is widely accepted that amino acids are stored in vacuoles, whereas phenylpropanoid pathway enzymes are mainly localized in the cytosol (Achnine *et al*., 2004; Hildebrandt *et al*., 2015; Shimada *et al*., 2018). It is unclear whether vacuole-stored Phe is exported back into the cytosol and utilized to boost phenylpropanoid production during HL exposure. Understanding Phe homeostasis at the subcellular resolution will provide fundamental insights into how Phe partitioning and inter-organellar transport constrain or enable metabolic flux toward phenylpropanoids. Also, obtaining a more precise and quantitative flux map of the phenylpropanoid pathway during HL stress, for example, through isotope-based metabolic flux analysis, will be essential to resolve how increased Phe availability is redistributed among competing downstream branches of not only anthocyanin but also other phenylpropanoids, including lignin (Matsuda *et al*., 2003; Simpson *et al*., 2021; Koley *et al*., 2024; Dong *et al*., 2025). Given the well-established roles of phenylpropanoids and anthocyanins in photoprotection and antioxidative defense, another important unresolved question is how the enhanced flux from Phe into phenylpropanoids under HL stress affects other physiological parameters, particularly reactive oxygen species production and redox homeostasis (Mittler *et al*., 2022; Cerqueira *et al*., 2023).

### Phe metabolism against abiotic stress in plants

Unlike the highly inducible phenylpropanoid and anthocyanin pathways, our proteomics reveals that most proteins involved in Phe biosynthesis remain unaltered by 2-day HL treatment. Strikingly, Phe levels were unchanged in Col-0 but decreased in *pap1D* after HL exposure (**Figure 1D**). These results suggest that Phe levels are sufficient in wild type to sustain phenylpropanoid production under HL stress, whereas artificial enhancement of anthocyanin production in *pap1D* leads to a precursor shortage when metabolic demand exceeds its Phe-producing capacity. Some studies report transcriptional upregulation of the *DHS1* gene in response to various abiotic and biotic stresses, such as UV, wounding, and jasmonic acid treatment, to induce the production of aromatic defense compounds (Keith *et al*., 1991; Görlach *et al*., 1995; Devoto *et al*., 2005; Yokoyama, Kleven, *et al*., 2022). Consistently, Kusano et al. uncovered that UV stress upregulates genes involved in primary metabolism, including aromatic amino acid biosynthetic genes, thereby increasing Phe availability for phenylpropanoid production (Kusano *et al*., 2011). However, our proteome analysis detected limited impacts on the shikimate and Phe biosynthetic enzymes, suggesting that HL exposure may activate a distinct gene regulatory set compared with UV stress. Notably, HL treatment affected a larger number of proteins in *pap1D* than in Col-0 (**Figure 2A and 2B**), possibly reflecting secondary effects of anthocyanin overaccumulation. Our proteomics also identified only a slight increase in protein levels of EPSPS2, CM1, and ADT3 in *pap1D* (**Figure 2F**), which may stimulate Phe biosynthesis. Given the reduced Phe levels in *pap1D* after HL exposure, however, this upregulation was insufficient to compensate for the Phe precursor shortage during HL stress. In contrast to the relatively unresponsive proteome of Phe biosynthetic enzymes, the levels of numerous enzymes in phenylpropanoid and anthocyanin pathways, including PAL1 and PAL2, which are inducible by environmental stimuli (Raes *et al*., 2003; Rohde *et al*., 2004; Barros and Dixon, 2020), were drastically elevated after HL exposure. Our proteomics data confirm that Phe biosynthesis is less responsive to HL exposure than Phe-derived specialized metabolism.

Why is Phe biosynthesis more stable in response to environmental changes? Here, we discuss three possibilities. First, Phe biosynthesis may be maintained stably to avoid perturbing core biological functions, such as growth and protein synthesis, which generally show limited responsiveness to environmental fluctuations (Son and Park, 2023; Coates *et al*., 2025). Second, because aromatic amino acids are more carbon- and energy-intensive than the other simpler amino acids (Arnold and Nikoloski, 2014), plants may produce and maintain Phe only to a level sufficient for short-term environmental changes, thereby minimizing unnecessary carbon and energy consumption. Third, limited Phe supply may act as a metabolic "brake" to prevent overproduction of anti-stress phenylpropanoids, particularly after stress conditions have subsided. Addressing these possibilities will require further physiological characterization of the *sota* mutants, such as application of longer and/or combinational stresses, to uncover the potential negative consequences of Phe overaccumulation.

### Lessons for plant metabolic engineering

Enhancing precursor availability is a common strategy to boost the production of target compounds in metabolic engineering. Recent efforts to study the diversification of plant aromatic amino acid metabolism have identified numerous lineage-specific enzyme isoforms that boost aromatic amino acid production, thereby enabling their use as a promising genetic toolbox to increase precursor availability (Maeda, 2019b; Yokoyama, 2024). Unlike microbes, however, this approach does not always result in substantial accumulation of target compounds in plants, owing to more complex, multilayered regulatory mechanisms at the interface between precursor supply and downstream pathways of primary and specialized metabolism (Maeda, 2019b; Yokoyama, 2024). To overcome this challenge, identifying metabolic bottlenecks at the precursor-to-product interface should be a key step to maximize the impact of enhanced precursor availability. A recent study with the focus on Tyr and betalains, Tyr-derived natural red pigments (Polturak and Aharoni, 2018), underscores the importance of the “push-pull” strategy for efficient plant metabolic engineering (Kacser and Acerenza, 1993; Jung and Maeda, 2024). This study demonstrates that combining enhanced Tyr supply (“push”) with effective downstream utilization of the precursor (“pull”) by relieving the bottleneck at the L-DOPA 4,5-dioxygenase step synergistically boosts heterologous betalain production, highlighting that increased precursor availability alone is insufficient without efficient metabolic draw through the pathway (Jung and Maeda, 2024). This lesson is directly applicable to our study, in which “increased Phe precursor supply conferred by the *sota* mutations” and “enhanced Phe utilization through anthocyanin production under HL exposure” can be classified as the “push” and “pull” components of the strategy, respectively, demonstrating the transferability of the "push and pull" strategy for plant metabolic engineering of Phe-derived high-value natural products. An important next step will be to identify which combinations of “push” and “pull” strategies most effectively maximize metabolic output, particularly because HL stress is neither an ideal nor a sustainable “pull” treatment and instead poses a potential threat to multiple aspects of plant physiology (Dixon and Paiva, 1995; Gould, 2004; Araguirang and Richter, 2022). In this context, identifying optimal mutation types and/or strategically selecting appropriate gene sources from species with naturally high metabolic flux toward the target metabolites will be critical for the rational design of robust and scalable plant metabolic engineering of phenylpropanoid compounds.

### Experimental Procedures

#### Plant growth and high-light treatment

*Arabidopsis thaliana* plants were sown directly on soil in 6-cm pots, and grown under a short-day photoperiod in a climate-controlled chamber (8-h light, 100 µmol m^-2^ s^-1^, day/night temperature of 20°C/16°C and humidity 60%/75%). For HL treatment, plants were transferred to another growth chamber and exposed to continuous HL stress (600 µmol m^-2^ s^-1^, 20°C, 60% humidity). The positions of the pots were regularly changed to minimize positional effects.

#### Generation of *sota pap1D* double mutants

The *pap1D* mutant was crossed with either *sotaB4* or *sotaA4* to obtain their F1 seeds. After the self-pollination of F1 plants, approximately 100 F2 plants were grown for genotyping. Homozygous *pap1D sota* double mutants were isolated and subsequently self-pollinated. The resulting F3 plants were genotyped again to confirm the presence of homozygous mutations and then used for phenotypic analyses. For the *sota* genotyping, derived Cleaved Amplified Polymorphic Sequences (dCAPS) genotyping was conducted to detect single-nucleotide polymorphisms as previously developed (Yokoyama, de Oliveira, *et al*., 2022). The *DHS1* and *DHS2* sequences were amplified using EconoTaq PLUS green 2x master mix (Lucigen) in a 20-μl reaction containing ∼10 ng of genomic DNA and 0.5 μM of each primer for *sotaB4* and *sotaA4*, respectively (**Supplemental Table 1**). The PCR products carrying the wild-type *DHS1* and *DHS2* sequence were digested with GsuI into 156 + 73 bp and 209 + 58 bp fragments, while GsuI did not cut the *sotaB4* or *sotaA4* mutated sequences, respectively. The digestion was confirmed on 4% 1×tris-borate EDTA (TBE)-agarose gel electrophoresis. For the *pap1D* genotyping, three primers listed in Supplemental Table 1 were mixed with EconoTaq PLUS green 2×master mix to amplify the wild-type and *pap1D* sequences with expected sizes of 646 and 1458 bp, respectively.

#### Sample harvesting and metabolite extraction

All fully expanded leaves of 5-6-week-old Arabidopsis plants were pooled from multiple plants at the same developmental stages, frozen in liquid nitrogen, ground with metal beads in a plastic bottle, and stored at -80°C prior to further analysis. For each analysis, 30-100 mg of each frozen sample was aliquoted from the frozen stocks and further analyzed as one biological replicate. Metabolite extraction was carried out as previously described (Yokoyama *et al*., 2021), with some modifications. Briefly, the frozen samples were vigorously vortexed for 10 min at room temperature in 800 µL of extraction buffer (v/v 2:1 of methanol and chloroform with 2 µg/mL isovitexin for peak normalization during LC-MS data analysis), followed by the addition of 600 µL of H_2_O and then 250 µL of chloroform. After another round of vortexing, the polar phase containing soluble metabolites was separated by centrifugation, aliquoted into three new tubes (300 µL each) designated for LC-MS analysis, anthocyanin quantification, and backup, then dried down and stored at -20 °C until further use.

### Anthocyanin quantification

For absorption-based anthocyanin quantification, the dried polar phase was dissolved in 500 µL of water. After adding 5 µL of 5 N HCl for acidification, the absorbance was measured at 530 and 657 nm using a microplate reader to calculate anthocyanin content using the formula A530 −0.25 × A657 (Mancinelli, 1990).

### LC-MS metabolomics

The dried soluble metabolite samples were dissolved in 100 µL of 80% methanol and analyzed on a Thermo Q Exactive Focus LC-MS platform equipped with a C18 column (40 °C; 400 µL min^-1^). Samples were run under a water/acetonitrile gradient containing 0.1% formic acid, and mass spectra were acquired in full-scan mode (*m*/*z* 100-1500) using data-independent acquisition (HCD 30 eV, positive ion mode). For metabolite identification, pooled quality-control samples were analyzed under both DIA and DDA modes. LC-MS full-scan data processing was performed using MS Refiner (Expressionist 14.0). Processing steps included retention-time alignment, peak detection based on mass reoccurrence and peak shape, noise filtering using predefined thresholds, isotope grouping, and removal of singleton features. Compound annotations were confirmed using authentic standards referenced in the Gold database. Multivariate statistical analyses, including principal component analysis and hierarchical clustering (heatmaps), were conducted to assess variation in metabolite profiles (Tiozon *et al*., 2023), using R version 2024.12.1+563.

### Preparation of total proteome and library

Frozen and ground Arabidopsis samples (30-60 mg; four biological replicates) were dissolved in 250 µL of extraction buffer (8M urea, 5 mM dithiothreitol) and incubated for 30 min with shaking, after which cell debris was removed by centrifugation. Protein concentration was determined using Pierce 660 nm protein assay. Following the sample alkylation using chloroacetamide (CAA) (550 mM stock, 14 mM final concentration) for 30 min at room temperature in the dark, aliquots corresponding to 50 µg total protein were subjected to in-solution digestion. In brief, the samples were diluted with 100 µL of a buffer containing 100mM Tris, pH 8.5, 1mM CaCl_2_, and 0.5 µg Lys-C (WAKO) was then added. After incubating the samples for 3h at 37 °C with shaking, 250 µL of a buffer containing 100mM Tris, pH 8.5 and 0.5 µg trypsin was added, followed by overnight incubation at 37 °C with shaking. The resulting samples were acidified with 15 µL of 20% trifluoroacetic acid (TFA) and split up for total proteome and library samples. For the library samples, 16 µL of each condition per replicate was mixed and submitted to SCX fractionation. To this end, StageTips were prepared using 6 layers of SPE disk (Empore Cation 2251 material) activated with acetonitrile and 1% TFA (100 µL each) and washed with buffer A (water, 0.2% TFA) (100 µL) by spinning 5 min (1500×*g*). The samples were then acidified to 1% TFA, loaded by centrifugation (10 min, 800×*g*) and washed with 100 µL of buffer A again. Fractionation was carried out using an ammonium acetate gradient in the 20% acetonitrile and 0.5 % formic acid (ACN and FA, respectively) solution starting from 25 mM to 500 mM for 9 fractions and two final elution steps using 1% ammonium hydroxide in 80% ACN and finally 5% ammonium hydroxide in 80% ACN. All fractions were eluted by centrifugation (5 min, 500×*g*) using 2 × 30 µL eluent. The fractions were dried and taken up in 10 µL A* buffer (2% ACN, 0.1% TFA). For measurement library samples were diluted to 0.2 µg/µL. Total proteome samples were desalted with C18 Empore disk membranes according to the StageTip protocol (Rappsilber *et al*., 2003). The eluted peptides were dried and then taken up in 10 µL A* buffer. Peptide concentration was determined by Nanodrop and samples were diluted at the 1:10 ratio with A* for measurement.

### LC-MS/MS data acquisition

Total proteome and fractionated (library) samples were analyzed using an Ultimate 3000 RSLC nano (Thermo Fisher) coupled to an Orbitrap Exploris 480 mass spectrometer equipped with a FAIMS Pro interface for Field asymmetric ion mobility separation (Thermo Fisher). Peptides were pre-concentrated on an Acclaim PepMap 100 pre-column (75 µM x 2 cm, C18, 3 µM, 100 Å, Thermo Fisher) using the loading pump and buffer A** (water, 0.1% TFA) with a flow of 7 µL/min for 5 min. Peptides were separated on 16 cm frit-less silica emitters (New Objective, 75 µm inner diameter), packed in-house with reversed-phase ReproSil-Pur C18 AQ 1.9 µm resin (Dr. Maisch). Peptides were loaded on the column and eluted for 130 min using a segmented linear gradient of 5% to 95% solvent B (0 min : 5%B; 0-5 min -> 5%B; 5-65 min -> 20%B; 65-90 min ->35%B; 90-100 min -> 55%; 100-105 min ->95%, 105-115 min ->95%, 115-115.1 min -> 5%, 115.1-130 min ->5%) (solvent A 0% ACN, 0.1% FA; solvent B 80% ACN, 0.1%FA) at a flow rate of 300 nL/min. Mass spectra were acquired in data-dependent acquisition mode with a TOP_S method using a cycle time of 2 seconds. For Field asymmetric ion mobility separation (FAIMS) two compensation voltages (-45 and -60) were applied. The cycle time for each experiment was set to 1 second. MS spectra were acquired in the Orbitrap analyzer with a mass range of 320-1200 *m*/*z* at a resolution of 60,000 FWHM and a normalized AGC target of 300%. Precursors were filtered using the MIPS option (MIPS mode = peptide), with the intensity threshold set to 5000, and were selected with an isolation window of 1.6 *m*/*z*. HCD fragmentation was performed at a normalized collision energy of 30%. MS/MS spectra were acquired with a target value of 75% ions at a resolution of 15,000 FWHM, at an automated injection time and a fixed first mass of *m*/*z* 100. Peptides with a charge of +1, greater than 6, or with unassigned charge state were excluded from fragmentation for MS^2^, dynamic exclusion for 40s prevented repeated selection of precursors.

### Proteomics data analysis

Raw data were processed using MaxQuant software (version 1.6.3.4, http://www.maxquant.org/) (Cox and Mann, 2008) with label-free quantification (LFQ) and iBAQ enabled (Tyanova *et al*., 2016). Library samples and DDA samples were grouped into separate parameter groups. In the group specific parameters, in the Misc. setting the Match type for library samples was set to “match from” and for DDA to “match from and to”. MS/MS spectra were searched by the Andromeda search engine against a combined database containing the sequences from *A. thaliana*(TAIR10_pep_20101214; ftp://ftp.arabidopsis.org/home/tair/Proteins/TAIR10_protein_lists/), and sequences of 248 common contaminant proteins and decoy sequences. Trypsin specificity was required with a maximum of two missed cleavages allowed. Minimal peptide length was set to seven amino acids. Carbamidomethylation of cysteine residues was set as fixed, oxidation of methionine and protein N-terminal acetylation as variable modifications. The match between runs option was enabled. Peptide-spectrum-matches and proteins were retained if they were below a false discovery rate of 1% in both cases.

Statistical analysis of the MaxLFQ values was carried out using Perseus (version 1.5.8.5, http://www.maxquant.org/). Quantified proteins were filtered for reverse hits and hits “identified by site” and MaxLFQ values were log_2_ transformed. After grouping samples by condition only those proteins were retained for the subsequent analysis that had three valid values in one of the conditions. Two-sample *t*-tests were performed using a permutation-based FDR of 5%. Alternatively, quantified proteins were grouped by condition and only those hits were retained that had 4 valid values in one of the conditions. Missing values were imputed from a normal distribution (1.8 downshift, separately for each column).

Differential protein abundance analyses and functional enrichment were performed in R (Version 2024.12.1). Gene Ontology (GO), Gene Set Enrichment Analysis (GSEA), and KEGG pathway enrichment were conducted using Bioconductor packages, with protein identifiers converted to gene-level annotations prior to analysis. GO term enrichment (biological process, molecular function, and cellular component) was carried out using clusterProfiler, applying a hypergeometric test with Benjamini-Hochberg correction to control the false discovery rate (FDR). GSEA was performed on ranked protein lists based on log2 fold change using clusterProfiler::GSEA, with significance determined at FDR < 0.05. KEGG pathway enrichment was conducted using enrichKEGG, with organism-specific annotations retrieved through KEGGREST. Only pathways passing an FDR threshold of 0.05 were retained. Differentially abundant proteins were visualized using volcano plots generated in ggplot2, plotting log2 fold change against -log10(adjusted *P*-value). Proteins meeting both the fold-change and FDR criteria were designated as significantly upregulated or downregulated. Hierarchical clustering and heatmap visualizations were produced using ComplexHeatmap with variance-stabilized or log-transformed abundance values. Principal component analysis (PCA) was performed to assess sample-level variance and proteome-wide responses.

### PAL enzymatic assay

PAL activity assay was conducted as previously described (Kim and Hwang, 2014), with some modifications. Total proteins were extracted from approximately 100 mg of frozen samples using 100 mM phosphate buffer (pH 6.0) containing 2 mM EDTA, 4 mM dithiothreitol, and 2% (w/w) polyvinylpyrrolidone. After centrifugation at 10,000×*g* for 10 min at 4°C, the protein content of the supernatant was quantified by Bradford assay (Bradford, 1976). The extract containing 10-50 μg of protein was incubated at 30°C for 60 min with 80 µL of 10 mM borate buffer (pH 8.7) and 20 µL of 20 mM Phe. The reaction was stopped by the addition of 10 µL of 5 M HCl and then centrifuged for 10 min at 10,000×*g* to pellet the denatured protein. Absorbance was measured at 290 nm before and after incubation. One unit (U) of PAL activity was defined as the amount of the enzyme that produced 1 nmol cinnamic acid per hour. Reactions without the substrate served as the blank control.

## Data availability

The metabolomics raw data used in Figure 1 and Figures 3 and 4 were deposited in the Zenodo repositories under DOI numbers 10.5281/zenodo.18236343 and 10.5281/zenodo.18273668, respectively. The proteomics data are available in PRIDE with the accession number PXD073142. The other data will be made available upon request.

## Acknowledgments

We thank Dr. Saleh Alseekh for his skillful assistance in LC-MS metabolome analysis. The original *pap1D* seeds were kindly provided by the group of Prof. Marisa Otegui (University of Wisconsin-Madison). This work was supported by Marie Skłodowska-Curie Actions Postdoctoral Fellowship to R.Y. The current research of R.Y. is supported through the Preparing Future Faculty (PFF) program at University of Missouri. S.C.S., A.H., and H.N. were supported by the Max Planck Society.

## Author Contributions

RY designed the research. RY performed most of the experiments. SCS, AH, and HN carried out proteomics. RNT and RY analyzed the data. HAM and ARF helped with the research. RNT and RY wrote the article. All authors read and approved the content of this article.

