## Supplemental Figures for "Enhanced phenylalanine biosynthesis amplifies light-stress-driven phenylpropanoid production in Arabidopsis"

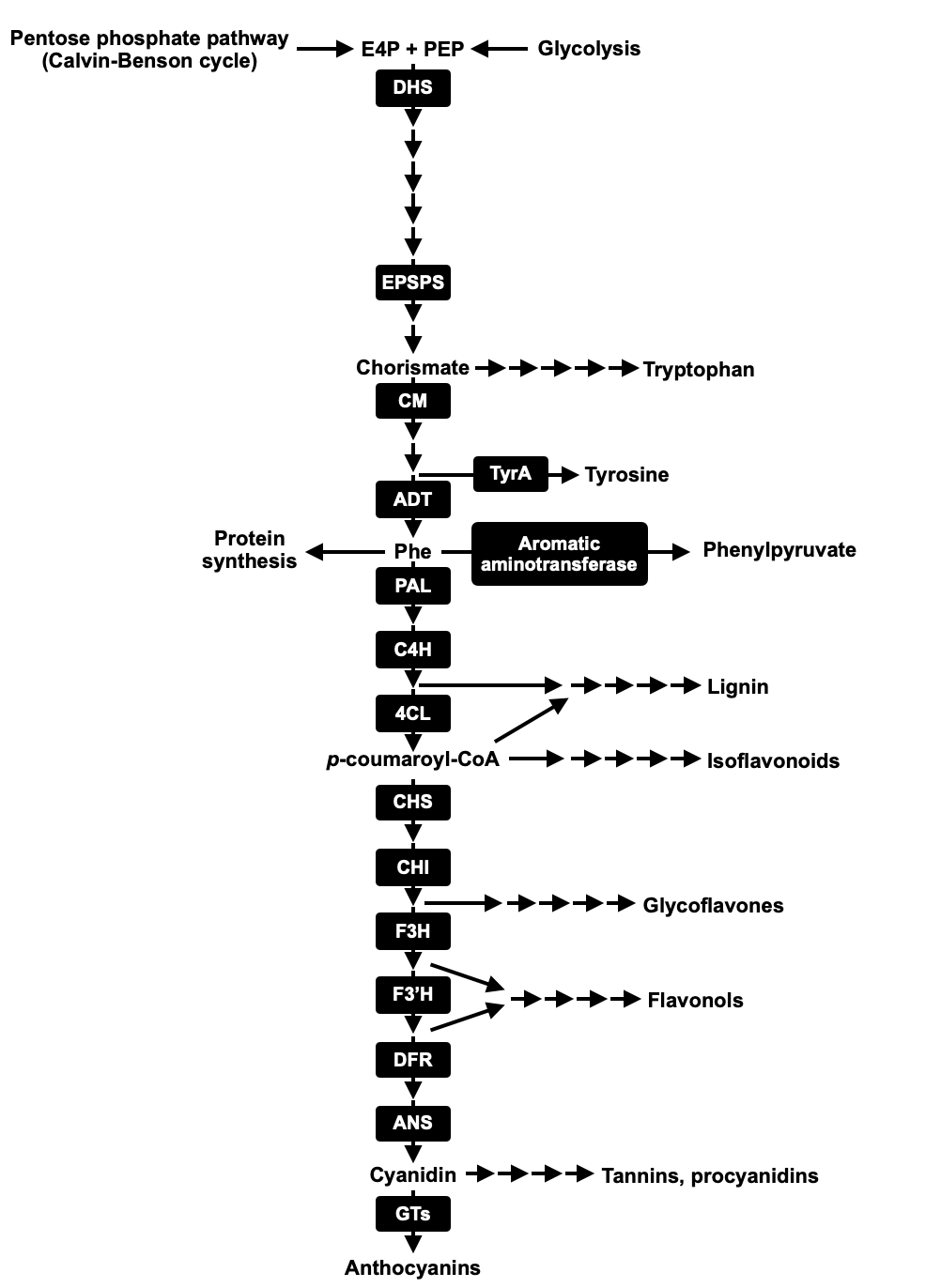

**Supplemental Figure 1. Pathway map of phenylalanine and phenylpropanoid biosynthesis.** DHS, 3-Deoxy-D-arabino-heptulosonate 7-phosphate (DAHP) synthase; EPSPS, Enolpyruvylshikimate 3-phosphate (EPSP) synthase; CM, Chorismate mutase; TyrA, Arogenate dehydrogenase; ADT, Arogenate dehydratase; PAL, Phenylalanine ammonia-lyase; C4H, Trans-cinnamate 4-hydroxylase; 4CL, 4-Coumarate:CoA ligase; CHS, Chalcone synthase; CHI, Chalcone isomerase; F3H, Flavanone 3-hydroxylase; F3'H, Flavonoid 3'-hydroxylase; DFR, Dihydroflavonol 4-reductase; ANS, Anthocyanin synthase; GTs, UDP-glucose:flavonoid glucosyltransferases.

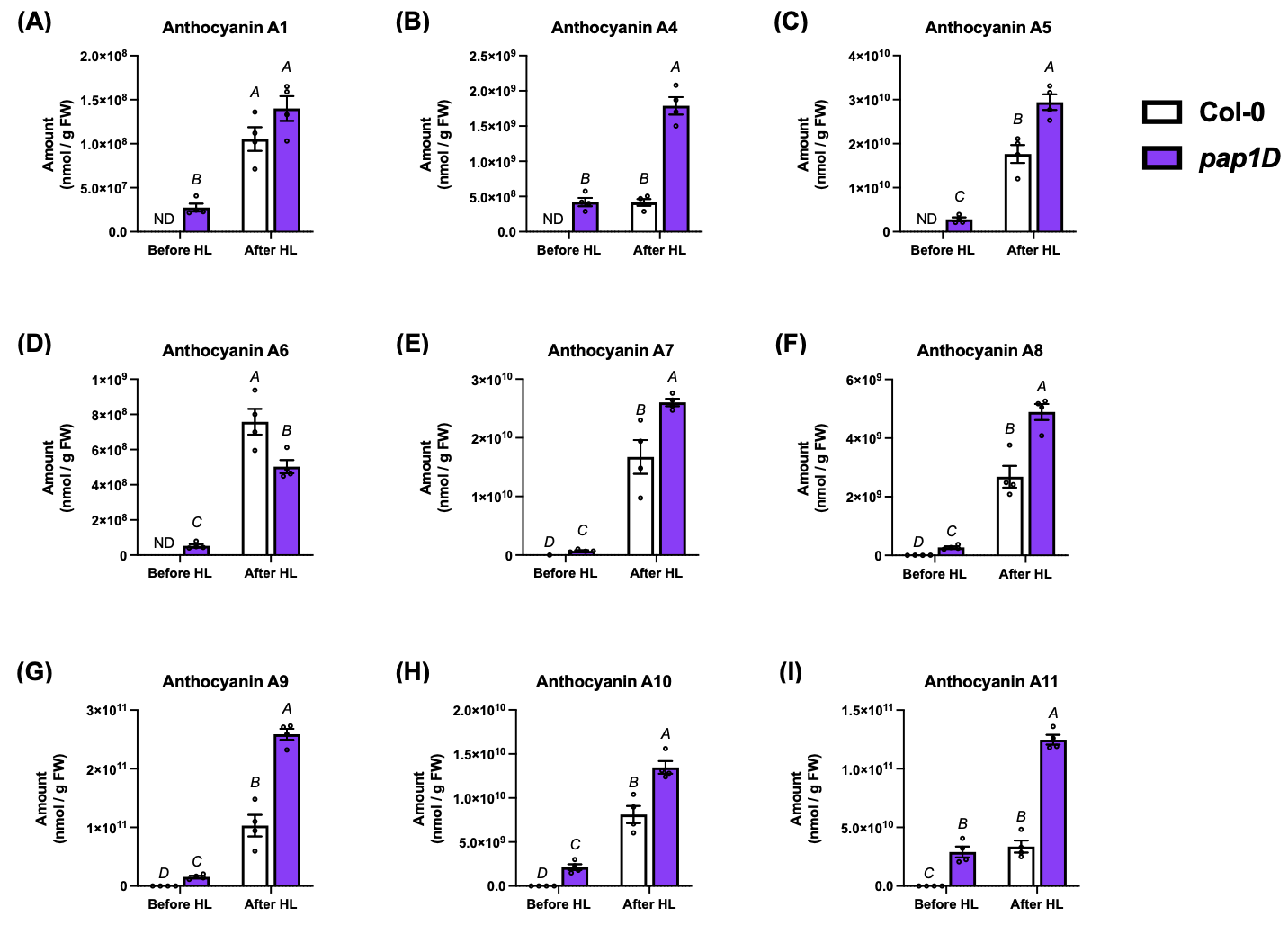

**Supplemental Figure 2. Accumulation of anthocyanin species before and after HL exposure.** Metabolite levels of anthocyanin A1 (A), A4 (B), A5 (C), A6 (D), A7 (E), A8 (F), A9 (G), A10 (H), and A11 (I) in Col-0 and *pap1D* before and after HL. Different letters indicate statistically significant differences (two-way ANOVA with Tukey-Kramer test, *P* < 0.05). Data are means ± SEM (*n* = 4). anthocyanin A1, Cyanidin 3-*O*-[2′′-*O*-(xylosyl) glucoside] 5-*O*-glucoside; anthocyanin A4, Cyanidin 3-*O*-[2′′-*O*-(2′′′-*O*-(sinapoyl) xylosyl) glucoside] 5-*O*-glucoside; anthocyanin A5, Cyanidin 3-*O*-[2′′-*O*-(xylosyl)-6′′-*O*-(*p*-coumaroyl) glucoside] 5-*O*-malonylglucoside; anthocyanin A6, Cyanidin 3-*O*-[2′′-*O*-(xylosyl)-6′′-*O*-(*p*-*O*-(glucosyl)-*p*-coumaroyl) glucoside] 5-*O*-glucoside; anthocyanin A7, Cyanidin 3-*O*-[2′′-*O*-(2′′′-*O*-(sinapoyl) xylosyl) 6′′-*O*-(*p*-coumaroyl) glucoside] 5-*O*-glucoside; anthocyanin A8, Cyanidin 3-*O*-[2′′-*O*-(xylosyl) 6′′-*O*-(*p*-*O*-(glucosyl) *p*-coumaroyl) glucoside] 5-*O*-[6′′′-*O*-(malonyl) glucoside]; anthocyanin A9, Cyanidin 3-*O*-[2′′-*O*-(2′′′-*O*-(sinapoyl) xylosyl) 6′′-*O*-(*p*-*O*-coumaroyl) glucoside] 5-*O*-[6′′′′-*O*-(malonyl) glucoside]; anthocyanin A10, Cyanidin 3-*O*-[2′′-*O*-(2′′′-*O*-(sinapoyl) xylosyl) 6′′-*O*-(*p*-*O*-(glucosyl) *p*-coumaroyl) glucoside] 5-*O*-glucoside; anthocyanin A11, Cyanidin 3-*O*-[2′′-*O*-(6′′′-*O*-(sinapoyl) xylosyl) 6′′-*O*-(*p*-*O*-(glucosyl)-*p*-coumaroyl) glucoside] 5-*O*-(6′′′′-*O*-malonyl) glucoside.

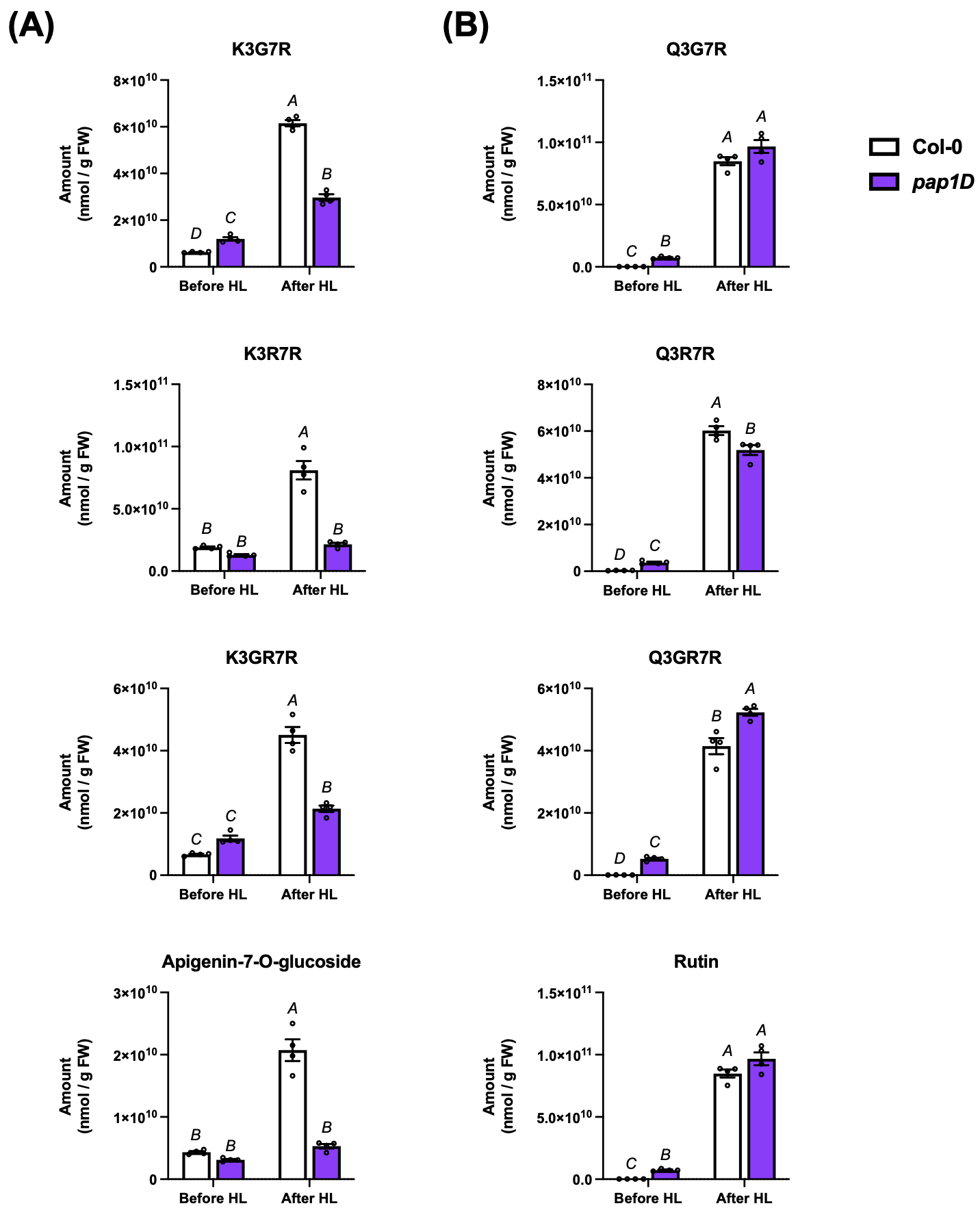

**Supplemental Figure 3. Accumulation of other major flavonoids before and after HL exposure.** Metabolite levels of kaempferol- or apigenin-conjugated flavonoids (A) and quercetin-conjugated flavonoids. Different letters indicate statistically significant differences (two-way ANOVA with Tukey-Kramer test, *P* < 0.05). Data are means ± SEM (*n* = 4). K3G7R, kaempferol-3-*O*-glucoside-7-*O*-rhamnoside; K3R7R, keampferol-3-*O*-rhamnoside-7-*O*-rhamnoside; K3GR7R, kaempferol-3-*O*-(2"-*O*-rhamnosyl)glucoside-7-*O*-rhamnoside; Q3G7R, quercetin-3-*O*-glucoside-7-*O*-rhamnoside; Q3R7R, quercetin-3-*O*-rhamoside-7-*O*-rhamnoside; Q3GR7R, quercetin-3-*O*-(2"-*O*-rhamnosyl)glucoside-7-*O*-rhamnoside; Rutin, quercetin-3-*O*-rutinoside.

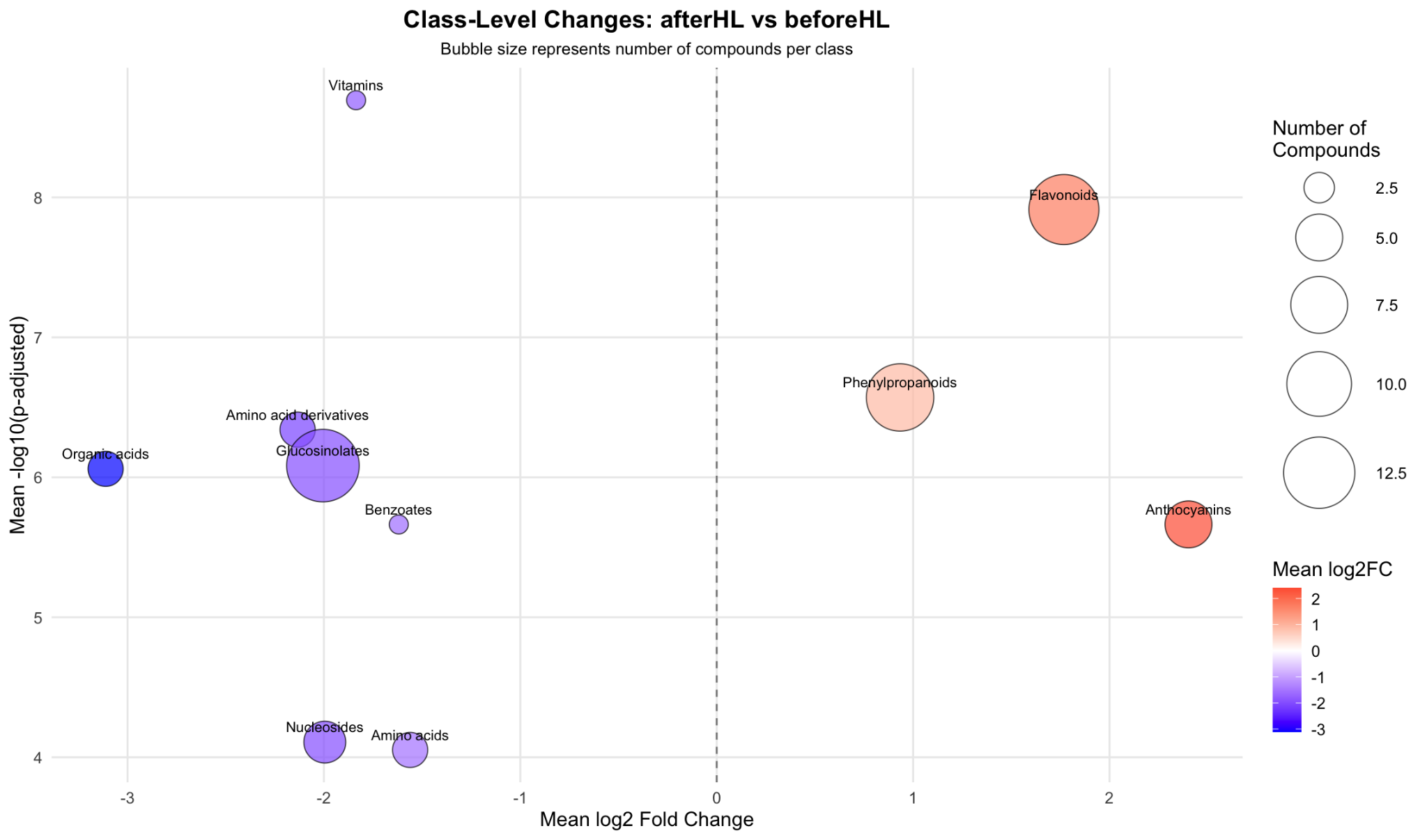

**Supplemental Figure 4.** Bubble plot summarizing class-level metabolite changes following HL treatment relative to control conditions. Each bubble represents a metabolite class, positioned according to the mean log₂ fold change (after HL vs before HL) on the *x*-axis and the mean -log₁₀ adjusted *p*-value on the *y*-axis. Bubble size indicates the number of annotated compounds within each class, and color denotes the direction and magnitude of the mean log₂ fold change.

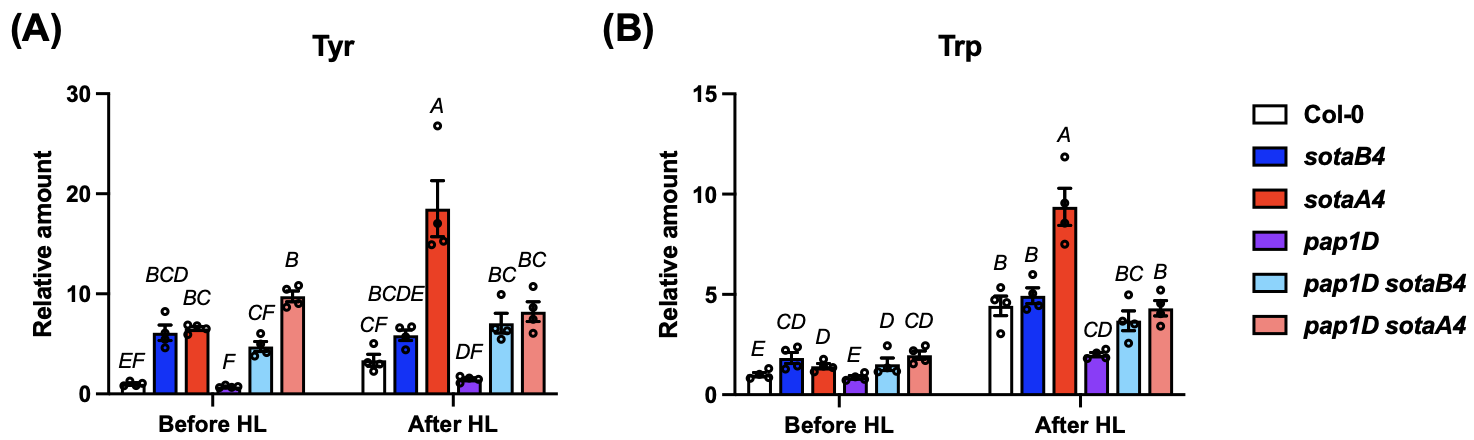

**Supplemental Figure 5. Accumulation of the other aromatic amino acids before and after HL exposure.** Metabolite levels of Tyr and Trp of Col-0, *sotaB4*, *sotaA4*, *pap1D*, *pap1D sotaB4*, and *pap1D sotaA4* before and after HL exposure. Different letters indicate statistically significant differences (two-way ANOVA with Tukey-Kramer test, *P* < 0.05). Data are means ± SEM (*n* = 4).

| Name | Sequence | For |
| --- | --- | --- |
| dCAPs_sotaB4_F | TTGAGGAGAAGGATGGAGTGA | dCAPS genotyping for sotaB4 |
| dCAPs_sotaB4_R | TCAGCTTGCTCACTTTGTTCA | dCAPS genotyping for sotaB4 |
| dCAPs_sotaA4_F | TTTTCAGGTGGGAAGAATGG | dCAPS genotyping for sotaA4 |
| dCAPs_sotaA4_R | TGTTGCGTGAAATCAAGGTT | dCAPS genotyping for sotaA4 |
| pap1D_LP | TAGCTCTAATGCTTGCTTACG | Genotyping for pap1D |
| pap1D_RP | ACTAAATCAGCCCAGGAAG | Genotyping for pap1D |
| pap1D_LBP | GTCCTGCCCGTCACCGAG | Genotyping for pap1D |

**Supplemental Table 1. The list of primers used in this study**

**Dataset S1.** Proteins commonly detected in Col-0 and *pap1D* under high-light conditions

**Dataset S2.** Differentially abundant proteins in wild-type (Col-0) under high-light stress (Day 0 vs Day 2)

**Dataset S3.** Differentially abundant proteins in *pap1D* under high-light stress (Day 0 vs Day 2)

**Dataset S4.** Differentially abundant proteins between wild-type (Col-0) and *pap1D* at Day 0 under standard light conditions

**Dataset S5.** Differentially abundant proteins between wild-type (Col-0) and *pap1D* after 2 days of high-light exposure

**Dataset S6.** Protein levels of enzymes in Phe and anthocyanin biosynthesis used in Figure 2F

**Dataset S7.** LC-MS metabolite annotation in positive mode

**Dataset S8.** LC-MS metabolite annotation in negative mode

**Dataset S9.** Metabolite levels obtained from untargeted metabolomics
